# Limited One-time Sampling Irregularity Map (LOTS-IM) for Automatic Unsupervised Assessment of White Matter Hyperintensities and Multiple Sclerosis Lesions in Structural Brain Magnetic Resonance Images

**DOI:** 10.1101/334292

**Authors:** Muhammad Febrian Rachmadi, Maria del C. Valdés-Hernández, Hongwei Li, Ricardo Guerrero, Rozanna Meijboom, Stewart Wiseman, Adam Waldman, Jianguo Zhang, Daniel Rueckert, Joanna Wardlaw, Taku Komura

## Abstract

We present the application of limited one-time sampling irregularity map (LOTS-IM): a fully automatic unsupervised approach to extract brain tissue irregularities in magnetic resonance images (MRI), for quantitatively assessing white matter hyperintensities (WMH) of presumed vascular origin, and multiple sclerosis (MS) lesions and their progression. LOTS-IM generates an irregularity map (IM) that represents all voxels as irregularity values with respect to the ones considered ”normal”. Unlike probability values, IM represents both regular and irregular regions in the brain based on the original MRI’s texture information. We evaluated and compared the use of IM for WMH and MS lesions segmentation on T2-FLAIR MRI with the *state-of-the-art* unsupervised lesions’ segmentation method, Lesion Growth Algorithm from the public toolbox Lesion Segmentation Toolbox (LST-LGA), with several well established conventional supervised machine learning schemes and with *state-of-the-art* supervised deep learning methods for WMH segmentation. In our experiments, LOTS-IM outperformed unsupervised method LST-LGA on WMH segmentation, both in performance and processing speed, thanks to the limited one-time sampling scheme and its implementation on GPU. Our method also outperformed supervised conventional machine learning algorithms (i.e., support vector machine (SVM) and random forest (RF)) and deep learning algorithms (i.e., deep Boltzmann machine (DBM) and convolutional encoder network (CEN)), while yielding comparable results to the convolutional neural network schemes that rank top of the algorithms developed up to date for this purpose (i.e., UResNet and UNet). LOTS-IM also performed well on MS lesions segmentation, performing similar to LST-LGA. On the other hand, the high sensitivity of IM on depicting signal change deems suitable for assessing MS progression, although care must be taken with signal changes not reflective of a true pathology.

## 1. Introduction

Stroke lesions, white matter hyperintensities (WMH) presumably of vascular origin and central nervous system’s lesions in multiple sclerosis (MS) similarly appear as hyperintense (bright) regions in T2-Fluid Attenuation Inversion Recovery (FLAIR) brain MRI. WMH of presumably vascular origin have been reported a predictor of stroke (Rensma et al., 2018) and are associated with cognitive decline (Pohjasvaara et al., 2000; del C. Valdés Hernández et al., 2013) and progression of dementia (Wardlaw et al., 2013). MS lesions described as T2-weighted hyperintensities of diameters of 3mm or more, disseminated in distinct anatomical locations within the central nervous system indicating a multifocal process, are part of the MS diagnostic criteria (Thompson et al., 2018). Interestingly, it has been reported that the extent of these namely T2-FLAIR hyperintensities not always predicts functional degeneration in MS, and that smaller T2-FLAIR hyperintensities could be disproportionally more damaging than larger lesions (Meier et al., 2007). Therefore the need to identify and characterise all breadth of lesions.

Due to their importance, there have been many studies proposing different approaches/methods for detecting and assessing WMH and MS lesions automatically. Hence, voxels of WMH and MS lesions in T2-FLAIR MRI have been identified and ”segmented” from the other ”normal” brain tissues either with the help of manually generated labels (supervised learning) or without the help of these manual labels (unsupervised learning).

Since the widespread use of deep neural network algorithms (i.e. referred to as ”deep learning”) in computer vision, these methods have become the *state-of-the-art* for detection/segmentation problems in brain MRI. For example, deep learning based architectures such as DeepMedic (Kamnitsas et al., 2017), UNet (Ronneberger et al., 2015) and UResNet (Guerrero et al., 2018) have outperformed conventional machine learning algorithms (e.g., support vector machine (SVM) and random forest (RF)) on automatic segmentation of WMH. However, as supervised methods, they are highly dependent on manual labels produced by experts (i.e., physicians) for training process. This dependency to expert’s opinion limits their applicability due to the expensiveness of manually generating WMH labels and the limited number of them. Furthermore, the quality of manual labels itself depends on and varies according to the expert’s skill. These intra/inter-observer inconsistencies can be quantified and reported, but they do not solve the problem. On the other hand, the more recent unsupervised deep learning methods based on generative adversarial networks (GAN) (Goodfellow et al., 2014), such as anomaly GAN (AnoGAN) (Schlegl et al., 2017) and adversarial auto-encoder (AAE) (Chen and Konukoglu, 2018)), need large number of both healthy and unhealthy data for adversarial training processes, usually not available or easily accessible.

On the other hand, conventional unsupervised segmentation methods, such as Lesion Growth Algorithm from Lesion Segmentation Tool toolbox (LST-LGA) (Schmidt et al., 2012) and Lesion-TOADS (Shiee et al., 2010), do not have the afore-mentioned dependencies to segment WMH and multiple sclerosis (MS) lesion in brain MRI. Hence, these methods have been tested in many studies and become the standard references to the other segmentation methods. Unfortunately, their performance is usually worse than that of supervised methods (Ithapu et al., 2014; Rachmadi et al., 2017a).

Recently, a new unsupervised segmentation method named irregularity age map (IAM) (Rachmadi et al., 2017b) and its faster version one-time sampling IAM (OTS-IAM) (Rachmadi et al., 2018c) have been proposed and reported to work better than the *state-of-the-art* unsupervised WMH segmentation method LST-LGA, the conventional machine-learning schemes SVM and RF, and some deep learning methods such as deep Boltzmann machine (DBM) (Salakhutdinov and Larochelle, 2010) and convolutional encoder network (CEN) (Brosch et al., 2016). IAM and OTS-IAM uniquely produce an irregularity map (IM) that has some advantage over deep learning’s probability map. Unlike a probability map, IM captures regular and irregular regions by retaining changes of the original T2-FLAIR intensities. This cannot be achieved with deep neural network algorithms, which are trained to reproduce manually generated binary masks (see Figure 2 and Figure 3). For example, the gradual changes of hyperintensities along the border of WMH, usually referred to as “penumbra” (Maillard et al., 2011), can be well represented in IM (see Figure 2). The penumbra of WMH has been subject of many studies in recent years, which debate criteria to correctly identify the WMH borders (Firbank et al., 2003; Jeerakathil et al., 2004; Hernández et al., 2010). Moreover, the penumbra of WMH itself is especially important for the study of WMH progression (Kapeller et al., 2003; Bendfeldt et al., 2009; Callisaya et al., 2013). It is also worth to mention that IM facilitates simulating the progression of WMH, as has been proposed previously (Rachmadi et al., 2018a).

**Figure 1:**
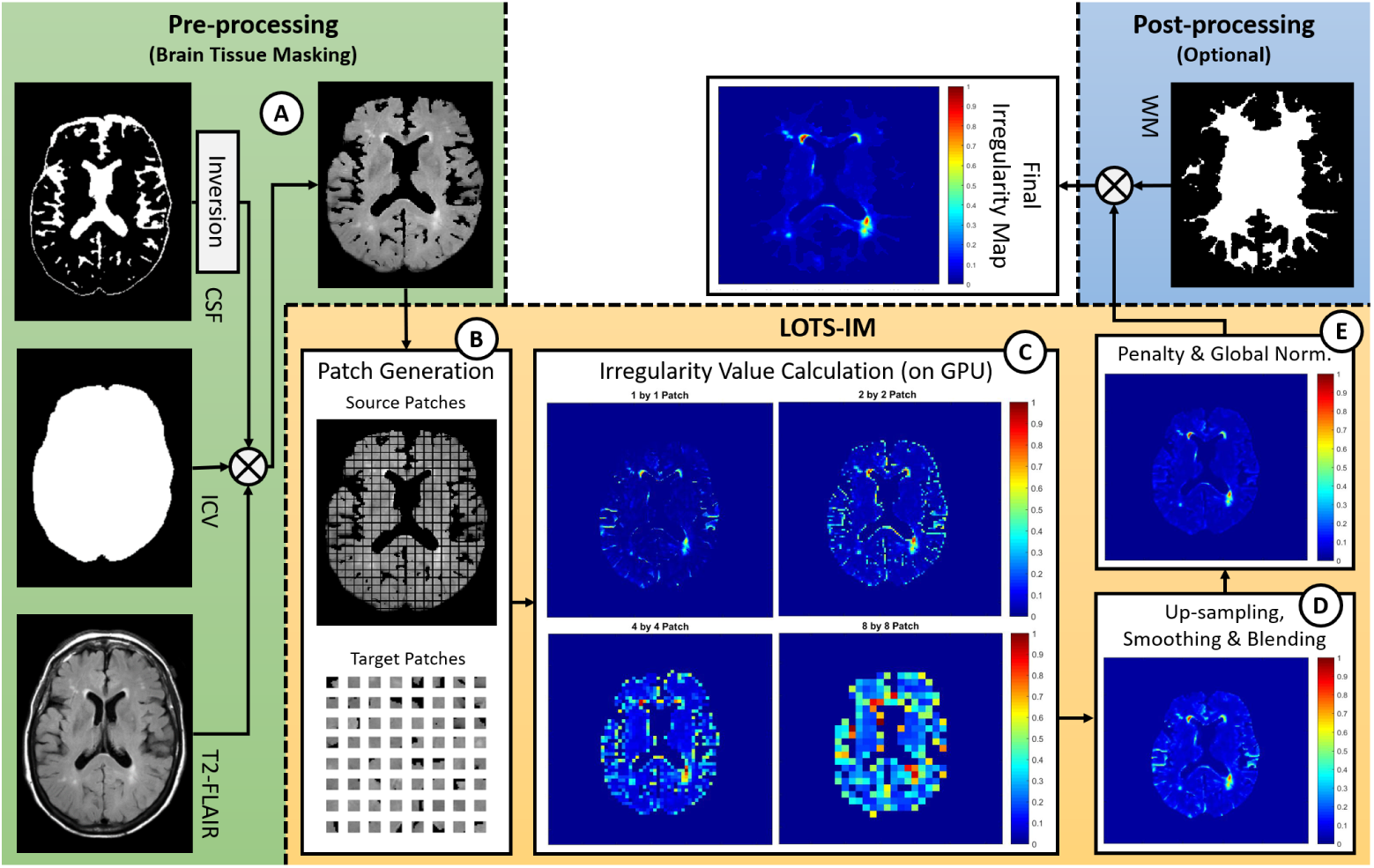
Flow of the proposed LOTS-IM. **1) Pre-processing**: brain tissue-only T2-FLAIR MRI 2D slices are generated from the original T2-FLAIR MRI and its corresponding brain masks (i.e., intracranial volume (ICV) and cerebrospinal fluid combined with pial regions (CSF)). **2) LOTS-IM**: the brain tissue-only T2-FLAIR MRI slice is processed through the LOTS-IM algorithm on GPU. **3) Post-processing**: final age map of the corresponding input MRI slice is produced after a post-processing step, which is optional.

**Figure 2:**
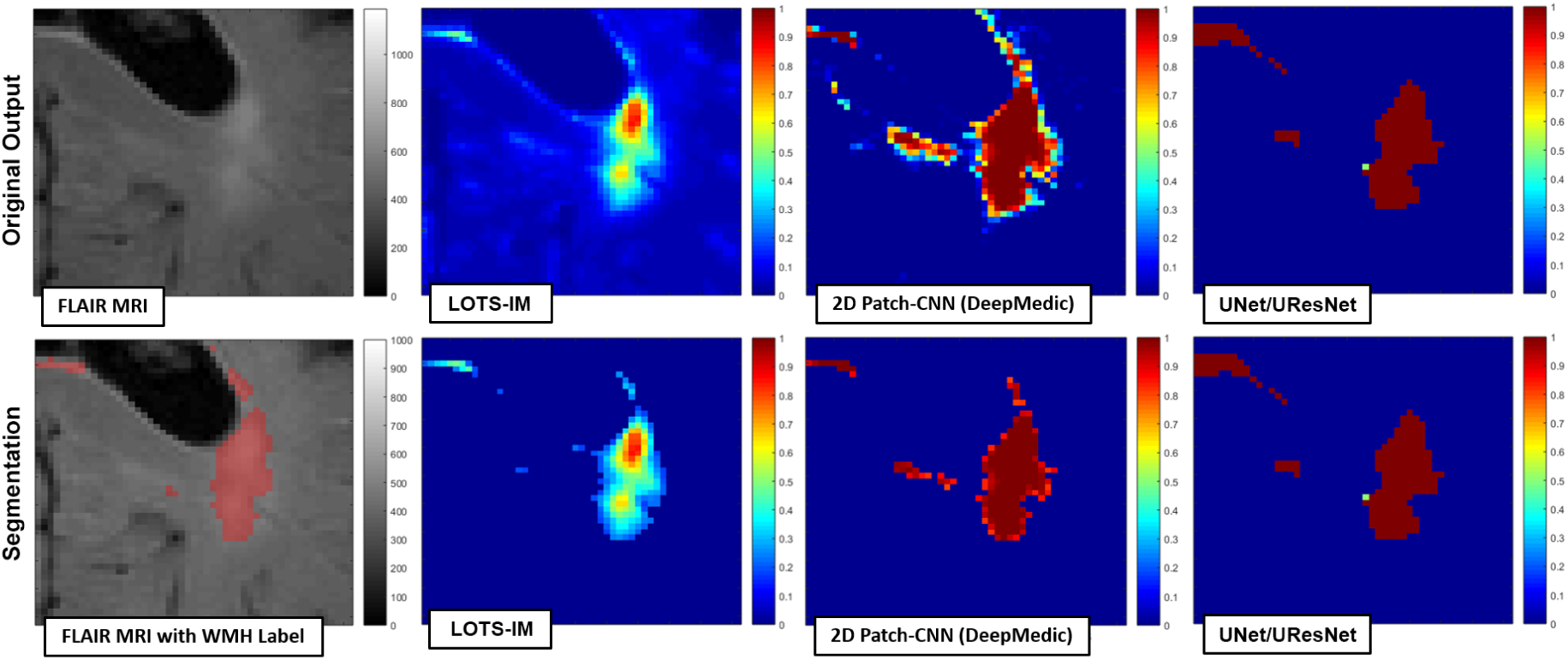
**Top:** Visualisation of original outputs produced by LOTS-IM (i.e., irregularity map) and other machine learning methods such as CNN, UNet and UResNet (i.e., probability maps). **Bottom:** Visualisation of WMH segmentation by cutting off the original values of irregularity/probability map. This figure shows that irregularity map not only nicely represents the penumbra of WMH by retaining the original textures but also is able to segment WMH by cutting off its values.

**Figure 3:**
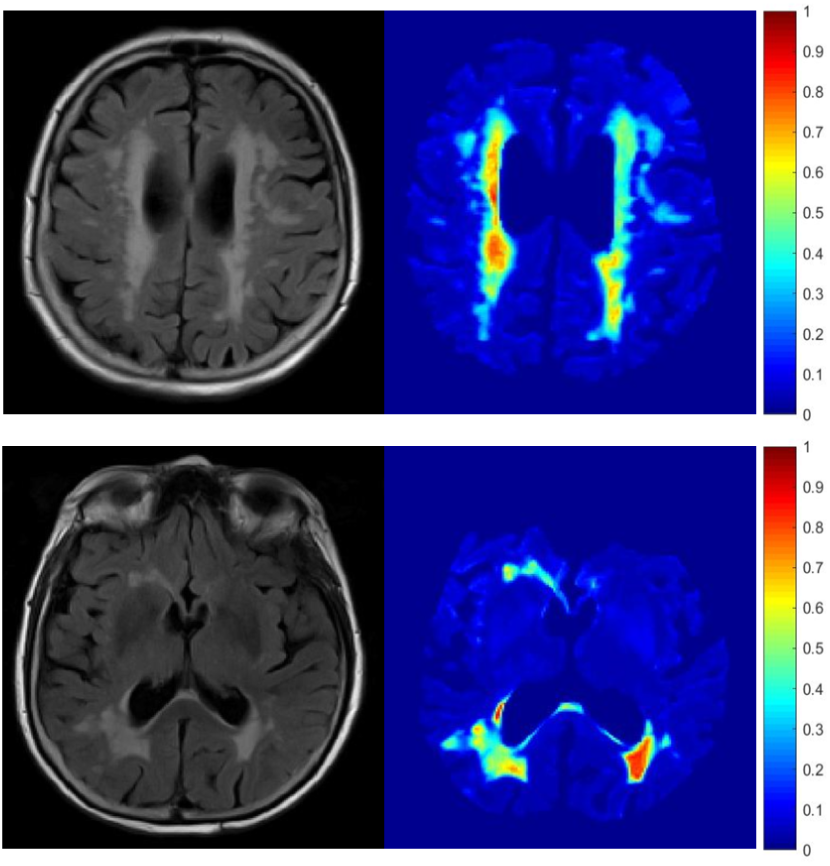
Large WMH visualised using irregularity map (IM) produced by the proposed method LOTS-IM. Note how both non-WMH and WMH regions, including the penumbra of WMH, are well represented by irregularity values.

While IAM and OTS-IAM have been tested in previous studies and show produced very good results in the segmentation of WMH in MRI scans from individuals with minor vascular pathology, they had one main limitation: their lengthy computing time. The most recent OTS-IAM takes 13 minutes (on GPU) to 174 minutes (on CPU) for processing a single MRI scan data of 256 × 256 × 35 voxels in average. The aforementioned computation times are not ideal especially if thousands of MRI are to be processed.

In this study, we proposed a new version of IAM namely limited one-time sample irregularity map (LOTS-IM) which greatly improves the processing time compared to IAM and OTS-IAM without having a perceivable quality degradation. This study also: 1) documents in more detail the generation of the irregularity map (IM) and the method’s performance (i.e. including limits of validity), not reported in previous studies, 2) describes and evaluates the generation of the internal parameters involved in the computation of the IM, and 3) evaluates the method’s performance in the segmentation of MS lesions.

## 2. Irregularity Map Method

The “irregularity age map” (IAM) for WMH assessment on brain MRI was originally proposed in (Rachmadi et al., 2017b), and it is based on a computer graphics method (Bellini et al., 2016) developed to detect aged/weathered regions in texture images. The term “age value” and “age map” were originally used by Bellini et al. (2016) for the 2D array of values between 0 and 1 denoting the weathered regions considered texture irregularities in natural images. In this study, we changed the term of age value and IAM to “irregularity value” and “irregularity map” as the concept of detecting “aged/weathered” textural regions no longer applies. In the irregularity map (IM), the closer the value to 1, the more probable a pixel/voxel belongs to a neighbourhood that has different texture from that considered “normal”.

After segmenting the regions of interest where the algorithm will work (e.g. brain tissue) using well established fully automatic computational methods (Step 1), IM is calculated from each structural MRI slice (i.e. preferably in axial or coronal orientation) by applying the following steps: patch generation (Step 2), irregularity value calculation (Step 3) and final irregularity map generation (Step 4). These four steps are schematically represented in Figure 1 and described in the rest of this section.

### 2.1. Brain tissue masking

For brain MRI scans, the brain tissue mask is necessary to exclude non-brain tissues which can represent “irregularities” *per se* (e.g., skull, cerebrospinal fluid, veins and meninges). In other words, we would like to compare brain tissue patches among themselves, not with patches from the skull, other extracranial tissues or fluid-filled cavities. For this purpose we use two binary masks: intracranial volume (ICV) and cerebrospinal fluid (CSF) masks, the latter containing also blood vessels and pial elements like venous sinuses and meninges. In our experiments, the ICV mask was generated by using optiBET (Lutkenhoff et al., 2014). However, several tools that produce accurate output exist and can be used for this purpose (e.g., bricBET^1^, freesurfer^2^). The CSF mask was generated by using a multispectral algorithm (Valdés Hernández et al., 2015). The brain tissue masking is schematically represented in Figure 1(A).

The pre-processing step before computing LOTS-IM only involves the generation of these two masks as per in the original IAM and in OTS-IAM (Rachmadi et al., 2017b, 2018b). Their subtraction generates the brain tissue mask, which is, then, multiplied by the FLAIR volume. This study also uses the normal appearing white matter (NAWM) mask in a post-processing step, as per OTS-IAM (Rachmadi et al., 2018b), to exclude brain areas in the cortex that could be identified as false positives. We generated NAWM masks using FSL-FAST (Zhang et al., 2001), but these can also be generated using other tools (e.g., freesurfer).

### 2.2. Patch generation

As per IAM Rachmadi et al. (2017b), LOTS-IM requires the generation of two sets of patches; non-overlapping grid patches called *source patches* and randomly-sampled patches called *target patches*, which can geometrically overlap each other (see Figure 1). In the IM computation, a source patch is used as reference to the underlying pixel/patch while a target patch is used to represent a sample of all possible image textures. A set of target patches is randomly sampled from the same image. Note that the distribution of randomly sampled target patches closely follows the underlying distribution of all target patches, i.e., brain tissues’ textures.

Source and target patches are used to calculate the irregularity value, where each of the source patches is compared with several randomly sampled target patches using a distance function (Bellini et al., 2016). This will be discussed in the next subsection. We use hierarchical subsets of four different sizes of source and target patches, which are 1 × 1, 2 × 2, 4 × 4 and 8 × 8 pixels. Thus, source and target patches are defined as 2D arrays. The patch generation process is schematically represented in Figure 1(B).

### 2.3. Irregularity value calculation

The *irregularity value* calculation is the core of the IM generation process. Let **s** be a source patch and **t** be a target patch, then the distance (*d*) between **s** and **t** is:

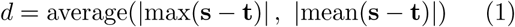

Based on Equation 1 above, the distance between source patch (**s**) and target patch (**t**) can be calculated by averaging the maximum difference and the mean difference between **s** and **t**. The difference between **s** and **t** is calculated by subtracting their intensities. The averaging of maximum and mean differences is applied to make the distance value robust against outliers. To capture the distribution of textures in an image (i.e., slice MRI), each source patch is compared against a set of target patches (e.g., 2,048 target patches in (Rachmadi et al., 2018c)) in which the same number of distance values are produced.

The *irregularity value* for a source patch can be calculated by sorting all distance values and averaging the 100 largest distance values of the whole set. The rationale is simple: the mean of the 100 largest distance values produced by an irregular source patch is still comparably higher than the one produced by a normal source patch. Also, mean is chosen as we are comparing irregularities with respect to the normal-appearing white matter, and normal tissue intensities are known to be normally distributed, although other descriptive statistics, such as percentiles, have been identified and used to discern degree of pathology (Dickie et al., 2015, 2014).

All irregularity values from all source patches are then mapped and normalised to real values between 0 and 1 to create the *irregularity map for one MRI slice* (see Figure 1(C)). Lastly, the IM is up-sampled to fit the original size of MRI slice (Rachmadi et al., 2018c) and smoothed using a Gaussian filter as per (Bellini et al., 2016).

### 2.4. Final irregularity map generation

The generation of the final IM consists of three sub-steps, which are a) *blending of the four irregularity maps produced in the irregularity value calculation step*, b) *penalty* and c) *global normalisation*.

*Blending of four irregularity maps* is performed by the following formulation:

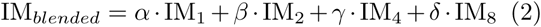

where *α* + *β* + *γ* + *δ* is equal to 1 and IM_1_, IM_2_, IM_4_ and IM_8_ are irregularity maps from 1 × 1,2 × 2, 4 × 4 and 8 × 8 pixels of source/target patches. Note that combining all information from patches of different sizes is performed to capture different levels of details, where smaller patches capture a more detailed information of the MRI’s intensity while bigger patches capture a bigger contextual information of the brain (Rachmadi et al., 2017b). Depiction of the blended IM can be seen in Figure 1(D).

The resulted (blended) irregularity map is then *penalised* using the formulation below:

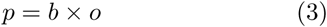

where *b* is the voxel from the blended irregularity map, *o* is voxel from the original MRI and *p* is the penalised voxel. Penalisation here is performed to eliminate artefacts usually caused by low quality ICV or CSF masks (Rachmadi et al., 2017b). Artefacts might be produced in previous step (Equation 1) when non-brain tissues represented as hypo-intensities in FLAIR MRI are unsuccessfully excluded by ICV and CSF masks. Note that Equation 1 cannot differentiate between hyper-intensities (i.e., bright voxels) and hypo-intensities (i.e., dark voxels) (Rachmadi et al., 2017b).

Lastly, all irregularity maps from different MRI slices are normalised together to produce values between 0 to 1 for each voxel to estimate “irregularity” with respect to the normal brain tissue across all slices. We name this normalisation procedure as *global normalisation*. Depiction of the resulted (i.e., penalised and globally normalised) irregularity map can be seen in Figure 1(E).

Some important notes on IM computation are: 1) source and target patches need to have the same size within the hierarchical framework, 2) the centre of source/target patches needs to be inside the brain and outside the CSF masks at the same time to be included in the irregularity value calculation and 3) the slice which does not provide any source patch (i.e where no brain tissue is observed) is skipped to accelerate computation.

## 3. Limited one-time sampling irregularity map (LOTS-IM)

As previously mentioned, while the original IAM has been reported to work well for WMH segmentation, its computation takes considerable time because it performs one target patch sampling for each source patch, selecting different target patches per source patch. For clarity, we named this scheme as *multiple-time sampling* (MTS) scheme. The MTS scheme is performed in the original IAM to satisfy the condition, stated in the original study (Bellini et al., 2016), that target patches should not be too close to the source patch (i.e., location based condition). Extra time in MTS to sample target patches for each source patch is, therefore, unavoidable under these premises.

To accelerate the overall IAM’s computation, we proposed and evaluated the one-time sampling (OTS) scheme, where target patches are randomly sampled only once for each MRI slice, hence abandoning the location based condition of the MTS (Rachmadi et al., 2018c). In other words, age values of all source patches from one slice were computed against one (i.e. the same) set of target patches. We named this combination of OTS scheme and IAM one-time sampling IAM (OTS-IAM).

In this study, we propose to limit the number of target patches to accelerate the overall computation, and we name our scheme limited one-time sampling IM (LOTS-IM). The original IAM, which runs on CPUs, uses an undefined large random number of target patches which could range from 10% to 75% of all possible target patches, depending on the size of the brain tissue in an MRI slice.

In the present study, six numbers of target patches are sampled and evaluated for the computation of LOTS-IM, which are 2048, 1024, 512, 256, 128 and 64. We also propose a more systematic way to calculate the irregularity value where the 1*/*8 largest distance values are used instead of a fixed number of 100. The ratio of the 1*/*8 largest distance values is used as it represents the second half of the third quartile (*Q*_3_) of the samples, i.e., outliers. Thus, the 256, 128, 64, 32, 16 and 8 largest distance values, deemed as outliers, are used to calculate irregularity values for 2048, 1024, 512, 256, 128 and 64 target patches respectively.

Smaller number of target patches in the LOTSIM scheme enables us to implement it on GPU to accelerate the computation. The limited number of samples in power-of-two is carefully chosen to ease GPU implementation, especially for GPU memory allocation.

## 4. Subjects and MRI Data

In this study, we use T2-Fluid Attenuation Inversion Recovery (T2-FLAIR) MRI from three different data sets to investigate the robustness and applicability of LOTS-IM. One data set comprises brain MRI data from mild cognitive impairment(MCI) and early Alzheimer’s disease (AD) patients while the other two are from multiple sclerosis (MS) patients.

The first data set used in this study contains 20 subjects from the Alzheimer’s Disease Neuroimaging Initiative (ADNI) (Mueller et al., 2005) database^3^. Each subject had three brain MRI scans obtained in three consecutive years. They were selected randomly and blind to any clinical, imaging or demographic information. All T2-FLAIR MRI sequences have the same dimension of 256 × 256 × 35 pixels where each voxel is 3.69*mm*^3^. Full data acquisition information appears described in (Rachmadi et al., 2018b). Ground truth was produced semi-automatically by an expert in medical image analysis using the region-growing algorithm in the Object Extractor tool in Analyze^*™*^ software guided by the co-registered T1-W and T2-W sequences. For more details on this data set, please see (Rachmadi et al., 2017a) and data-share page^4^. The investigators within the ADNI^5^ contributed to the design and implementation of ADNI and/or provided data but did not participate in the analysis or writing of this report.

The second data set contains brain MRI data from 30 MS patients with different loads of MS lesions, imaged in a 3T Siemens Magnetom Trio MR system at the University Medical Center Ljubljana (UMCL) (Lesjak et al., 2018). The data set is publicly available from ^6^ where co-registered T1-W, T2-W, T2-FLAIR and MS lesions ground truth are available. All MRI sequences for all subjects have the same dimension of 192 × 512 × 512 pixels where each voxel is 0.18*mm*^3^. Full data acquisition information can be looked at in (Lesjak et al., 2018).

The third data set is a set of longitudinal brain MRI data from 10 treatment-free multiple sclerosis (MS) patients, participants in the Future MS study: a multicentre study of this disease^7^. Images were acquired on a Verio 3T MRI scanner with a 20-channels head coil. Two experts visually assessed the images and identified the new lesions, enlarged lesions, and rated the disease progression in none, low, moderate or high. We compared the LOTS-IM output with the expert assessments and explored the applicability of this approach to studies of MS.

## 5. Other WMH Segmentation Methods used for evaluation

As LOTS-IM is an unsupervised method, we compare LOTS-IM’s performance with that from other unsupervised segmentation method: the lesion growth algorithm from the Lesion Segmentation Tool (LST-LGA) (Schmidt et al., 2012). The latter has been widely used as the main unsupervised reference standard method (Guerrero et al., 2018) for WMH and MS lesion segmentation, representing the *state-of-the-art* unsupervised WMH and MS lesions segmentation method. We used LST-LGA’s kappa value of 0.05 for both WMH and MS lesions segmentation, as in (Rachmadi et al., 2017a).

In the first data set (i.e. from the ADNI database), we also compare the performance of LOTS-IM with that from several supervised machine learning algorithms that have been previously tested and are commonly used for WMH segmentation. This comparison aims to give broader insight of LOTS-IM’s performance compared to other machine learning WMH segmentation methods, especially those using deep learning. The supervised methods evaluated in this study are Support Vector Machine (SVM), Random Forest (RF), Deep Boltzmann Machine (DBM), Convolutional Encoder Network (CEN), patch-based 2D CNN with global spatial information (DeepMedic-GSI-2D), patch-based 2D UResNet (Patch2D-UResNet) and patch-based 2D UNet (Patch2D-UNet). SVM and RF are chosen to represent conventional machine learning algorithms commonly used for WMH segmentation in many previous studies. DBM, CNN, DeepMedic and U-Net based methods are chosen to represent supervised deep learning methods commonly applied for WMH segmentation in recent years.

All supervised segmentation methods used in this study were trained and tested using 5-fold cross validation and evaluated on all 60 WMH labelled ADNI MRI scans. For clarity, we do not further elaborate on the implementation of the aforementioned algorithms. All of them were implemented as per previous studies: configurations for SVM, RF, DBM and CEN algorithms are described in detail in (Rachmadi et al., 2017a), whereas configurations and parameters for DeepMedic-GSI-2D, Patch2D-UResNet and Patch2D-UNet can be found in (Rachmadi et al., 2018b), (Guerrero et al., 2018) and (Li et al., 2018) respectively.

## 6. Evaluation Metrics

Dice similarity coefficient (DSC) (Dice, 1945), which measures similarity between ground truth and automatic segmentation results, is used here as the primary metric of evaluation. Higher DSC score means better performance, and the DSC score itself can be computed as follow:

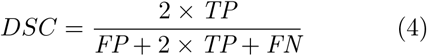

where *TP* is true positive, *FP* is false positive and *FN* is false negative.

Additional metrics positive predictive value (PPV), specificity (SPC) and true positive rate (TPR) are also calculated. Non-parametric Spearman’s correlation coefficient (Myers et al., 2010) is used to compute correlation between WMH volume produced by each segmentation method and visual ratings of WMH. Visual ratings of WMH are commonly used in clinical studies to describe and analyse severity of white matter disease (Scheltens et al., 1993). Correlation between visual ratings and volume of WMH is known to be high (Hernández et al., 2013). In this study, Fazekas’s (Fazekas et al., 1987) and Longstreth’s visual rating scales (Longstreth et al., 1996) are used for evaluation of each automatic method, as per (Rachmadi et al., 2017a).

Furthermore, two-sided Wilcoxon signed rank test is performed to see whether the difference between the performance of two algorithms is significant or not. The test produces two values; *p*-value and *h*. The latter shows the result of testing the null hypothesis that there is no significant difference of performance between the two algorithms (i.e., if h=1 the null hypothesis is rejected, and if h=0 the null hypothesis is not rejected). In this study, if *p* < 0.05 then the null hypothesis is rejected (i.e., *h* = 1).

## 7. Experiments and Results

This section is divided into two subsections. The first subsection presents the results from the MCI/AD patient (ADNI) data set, and the second subsection from the MS patients’ data sets. The ADNI data set was used not only to evaluate applicability of LOTS-IM for WMH segmentation but also to evaluate different aspects of this method (e.g., speed and its internal parameters of blending weights). The MS patients’ data sets were used to evaluate the applicability of LOTS-IM for MS lesion segmentation and progression analysis.

### 7.1. Results on MCI/AD (ADNI) Data Set

In this subsection, LOTS-IM is evaluated for WMH segmentation, longitudinal WMH assessment, and compared with supervised and unsupervised methods. Amongst the latter are the original IAM and OTS-IAM. In addition, in this subsection we evaluate LOTS-IM’s performance for scans with different WMH burden and analyse its speed, blending weights and random sampling.

#### 7.1.1. LOTS-IM for WMH segmentation

Table 1 shows the performance of all methods evaluated for WMH segmentation. Please note that the original IAM is listed as IAM-CPU. Please also note that different optimum thresholds (i.e. TRSH in Table 1) were used to produce the best WMH segmentation for each methods. The best values of DSC, PPV, SPC and TPR evaluation metrics are underlined.

**Table 1:**
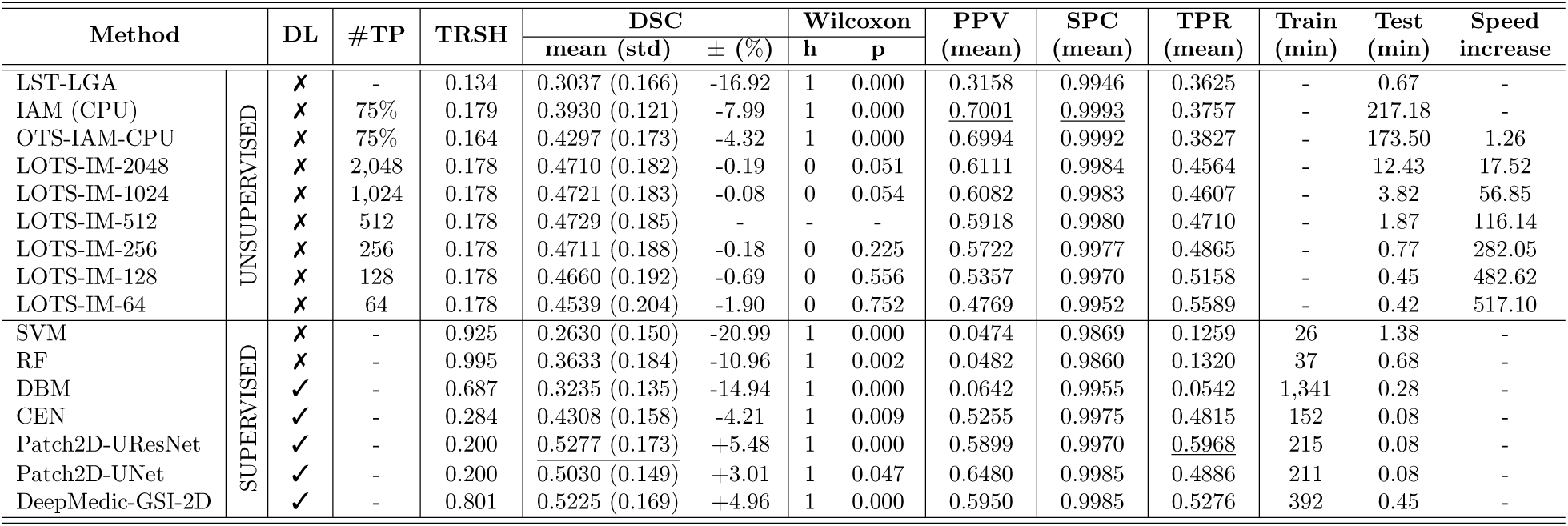
Experiment results of WMH segmentation based on Dice similarity coefficient (DSC), positive predictive value (PPV), specificity (SPC) and true positive rate (TPR). Best values for each metrics are <u>underlined</u>. Column “ ± (%)” shows relative performance difference (mean of DSC) to the LOTS-IM-512. Two-sided Wilcoxon signed rank test (with 5% significance level) is performed between LOTS-IM-512 and other methods to see whether the performance difference is significant or not. On the other hand, “Speed increase” is relative to IAM-CPU. **Abbreviations**: “DL” for deep learning method, “#TP” for number of target patches, “TRSH” for optimum threshold and “Train/Test” for training/testing time in minute (min).

From Table 1, we can see that the binary WMH segmentations produced by all IM method configurations (i.e., IAM, OTS-IAM and LOTS-IM methods) outperformed LST-LGA in mean DSC, PPV, SPC and TPR metrics. Especially for LOTS-IM-512, the best performer of all LOTS-IM methods, the performance differed up to 16.92% compared to LST-LGA. Furthermore, IAM/OTS-IAM/LOTS-IM not only outperformed LST-LGA but also conventional supervised machine learning algorithms (i.e., SVM and RF). Also, some LOTS-IM implementations outperformed supervised deep learning methods of DBM and CEN in DSC metric. Based on the two-sided Wilcoxon signed rank test, the performance of all LOTS-IM configurations were significantly different to LST-LGA, SVM, RF and DBM with *p <* 0.05.

Visual appearance of the irregularity map (IM) from LOTS-IM and probability maps produced by other segmentation methods such as DeepMedic-GSI-2D, and UNet/UResNet can be observed and compared in Figure 2. Figure 2 (top) shows that the IM produced by LOTS-IM retains the texture information of both non-WMH and WMH regions, including penumbra of WMH. IM can be used for WMH segmentation by thresholding its values, as shown in Figure 2 (bottom). Visualisation of the IM on a scan with large WMH load can be seen in Figure 3. Note how the penumbra of WMH is well represented in the IM in Figures 2 and 3. On the other hand, the probability maps produced by DeepMedic-GSI-2D and UNet/UResNet lack the ability to represent non-WMH regions and WMH penumbra.

Figure 4 shows the DSC performance curves of LOTS-IM and other WMH segmentation methods by cutting off the irregularity/probability values on different threshold values. LOTS-IM uses lower threshold values than the other methods to produce better WMH segmentation as the IM gives finer brain tissues details than the other methods (see Figure 2 and 3).

**Figure 4:**
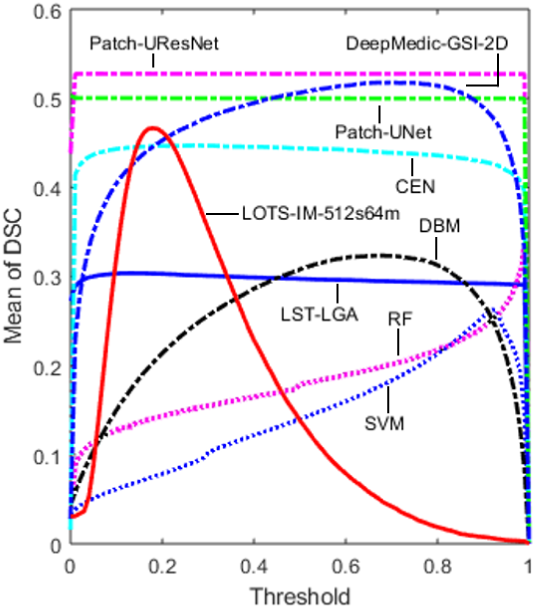
Mean of dice similarity coefficient (DSC) score for LST-LGA, SVM, RF, DBM, CEN, Patch2D-UResNet, Patch2D-UNet, DeepMedic-GSI-2D and LOTS-IM-512 in respect to all possible threshold values.

#### 7.1.2. LOTS-IM vs. IAM and OTS-IAM

Limited one-time sampling (LOTS) not only accelerated the computation time but also improved the overall performance, as shown in Table 1. Implementation of LOTS-IM on GPU increased the processing speed by 17 to 435 times with respect to the original IAM which was implemented on CPU. However, it is worth stressing that this increase in processing speed was not only due to the use of GPU instead of CPU, but also due to the limited number of target patch samples used in the the computation of LOTS-IM. Furthermore, one of the GPU implementations of LOTS-IM (i.e., LOTS-IM-64) ran faster than LST-LGA. Note that the testing time listed in Table 1 excludes registrations and the generation of other brain masks used either in pre-processing or post-processing steps. The increase in speed achieved by the GPU implementation of LOTS-IM shows the effectiveness of the proposed method in terms of computation time and overall performance.

#### 7.1.3. Speed vs. quality of LOTS-IM

The biggest contribution of this work is the increase in processing speed without compromising the quality of the results, achieved by LOTS-IM implemented on GPU, compared to the original IAM and OTS-IAM. The first iteration of IAM can only be run on CPU because it uses multiple-time sampling (MTS). OTS-IAM samples patches only once, but still uses a high number of target patches to compute the IM. Through this study, we show that using a limited number of target patches leads not only to faster computation but also to achieve small to none quality degradation.

The relation between speed and quality of the output (mean DSC) produced by IAM, OTS-IAM and all configurations of LOTS-IM is illustrated in Figure 5. Note that Figure 5 is extracted from Table 1. Also, it is worth mentioning that the use of more target patches in LOTS-IM produced better PPV and SPC evaluation metrics than when less target patches were used (i.e. also in LOTS-IM). The TPR metric, on the contrary, is better when less target patches are used compared to when the number of target patches is higher.

**Figure 5:**
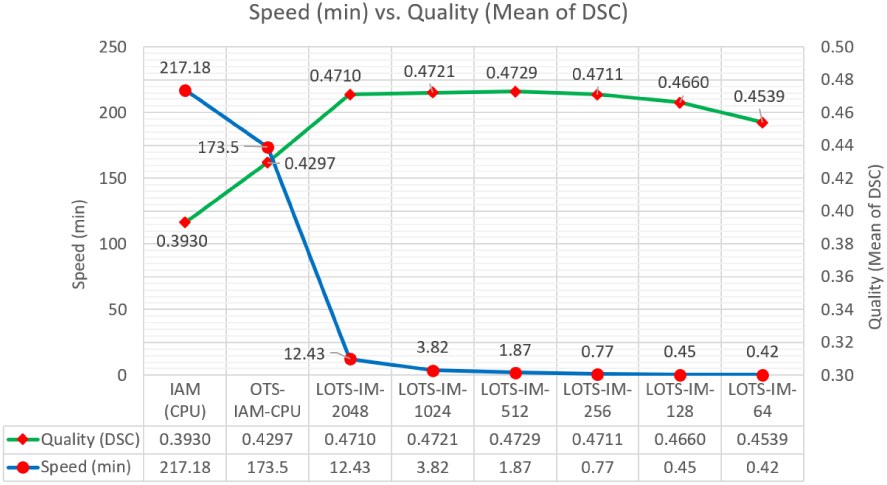
Speed (min) versus quality (mean of DSC) of different settings of LOTS-IM (extracted from Table 1). By implementing LOTS-IM on GPU and limiting the number of target patch samples, computation time and result’s quality are successfully improved and retained.

The two-sided Wilcoxon signed rank test also shows that there was no significant difference between LOTS-IM methods (i.e. *p* ≥ 0.05). Thus, LOTS-IM is more flexible than other methods in terms of speed as its computation speed can be adjusted as needed without compromising the output’s quality.

#### 7.1.4. Analysis on LOTS-IM’s blending weights

LOTS-IM has four internal parameters used to blend four irregularity maps, hierarchically produced by four different sizes of source/target patches, to generate the final IM (see Equation 2 in Section 2.4). In this experiment, different sets of blending weights in LOTS-IM’s computation were evaluated.

We tested 7 different sets of blending weights, which are listed in Table 2. The first 4 sets only use one of the irregularity maps (i.e., either the IM from 1 × 1, 2 × 2, 4 × 4 or 8 × 8 pixels). On the other hand, the last 3 sets blend all four irregularity maps with different blending weights. The effect of different sets of blending weights is illustrated in Figure 6.

**Table 2:** Mean and standard deviation of DSC produced by using different settings of blending weights. Plots corresponding to settings listed in this table can be seen in Figure 6. The LOTS-IM tested in this experiment is LOTS-IM-512.

| Name | Blending Weights |  |  |  | TRSH | DSC |  |
| --- | --- | --- | --- | --- | --- | --- | --- |
|  | 1x1 | 2x2 | 4x4 | 8x8 |  | mean | std |
| LST-LGA | - | - | - | - | 0.134 | 0.2936 | 0.1658 |
| IM-1000 | 1 | 0 | 0 | 0 | 0.128 | 0.4555 | 0.1774 |
| IM-0100 | 0 | 1 | 0 | 0 | 0.267 | 0.3995 | 0.1646 |
| IM-0010 | 0 | 0 | 1 | 0 | 0.376 | 0.3439 | 0.1627 |
| IM-0001 | 0 | 0 | 0 | 1 | 0.495 | 0.2594 | 0.1289 |
| IM-balanced | 0.25 | 0.25 | 0.25 | 0.25 | 0.287 | 0.4158 | 0.1754 |
| IM-4321 | 0.40 | 0.30 | 0.20 | 0.10 | 0.228 | 0.4486 | 0.1776 |
| IM-default | 0.75 | 0.19 | 0.05 | 0.01 | 0.179 | 0.4692 | 0.1820 |

**Figure 6:**
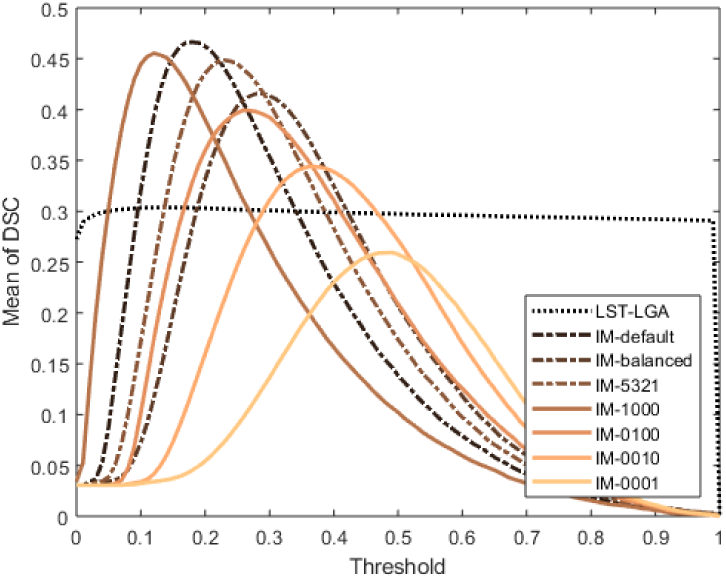
Curves of mean dice similarity coefficient (DSC) produced by using different settings of blending weights. LOTS-IM used in this experiment is LOTS-IM-512, and all weights are listed in Table 2.

From Figure 6 and Table 2, we can see that blending irregularity values from different irregularity maps produced better WMH segmentation results. Also, the IM produced by 1 × 1 pixels of source/target patches influences the WMH segmentation more than the others (i.e. those of dimensions 2 × 2, 4 × 4 and 8 × 8 pixels). The skewed blending weights of 0.75, 0.19, 0.05 and 0.01 produced the best DSC score. The skewed blending weights come from the ceiling operation of normalising by a power of two (i.e., 2^6^*/*85, 2^4^*/*85, 2^2^*/*85 and 2^0^*/*85 where 85 = 2^6^ + 2^4^ + 2^2^ + 2^0^). Based on the two-sided Wilcoxon signed rank test, the performances of the skewed blending weights to the IM produced by 1 × 1 pixels of source/target patches were significantly different to the others (i.e. *p <* 0.05). As the skewed blending weights of 0.75, 0.19, 0.05 and 0.01 produced the best DSC score in this experiment, we made this blending set to become the default blending set for the LOTS-IM. Also, note that this default blending set was used for all other experiments in Section 7.1 (i.e., Subsections 7.1.1, to 7.1.8) and Section 7.2 (i.e., Subsections 7.2.1 and 7.2.2).

Through this experiment, we see that it is necessary to consider not only the intensity of the individual pixels but also those from the group of pixels (textons, which convey the textural information). Combining irregularity maps produced by different sizes of non-overlapping sources is also similar to calculating IM using overlapping source patches. Furthermore, it is also useful to reduce pixellation or discretisation of IM due to voxel-wise computation. Nevertheless, this experiment shows that individual pixel intensities constitute the strongest feature for irregularity detection.

#### 7.1.5. WMH burden scalability test

In this experiment, all methods were evaluated to see their performances on segmenting WMH in MRI scans with different burden of WMH. The DSC metric is still used, but the data set is grouped into three different groups according to each patient’s WMH burden. These groups are listed in Table 3, and the results of this experiment can be seen in Figure 7 and Table 4. Please note that LOTS-IM is represented by LOTS-IM-512 as the best performer amongst the LOTS-IM methods (see Table 1).

**Table 3:** Three groups of MRI data based on WMH volume.

| No. | Group | WMH Vol. ($mm^3$) | # MRI Data |
| --- | --- | --- | --- |
| 1 | Small | WMH $\leq 4500$ | 27 |
| 2 | Medium | 4500 < WMH $\leq 13000$ | 25 |
| 3 | Large | WMH > 13000 | 8 |

**Table 4:**
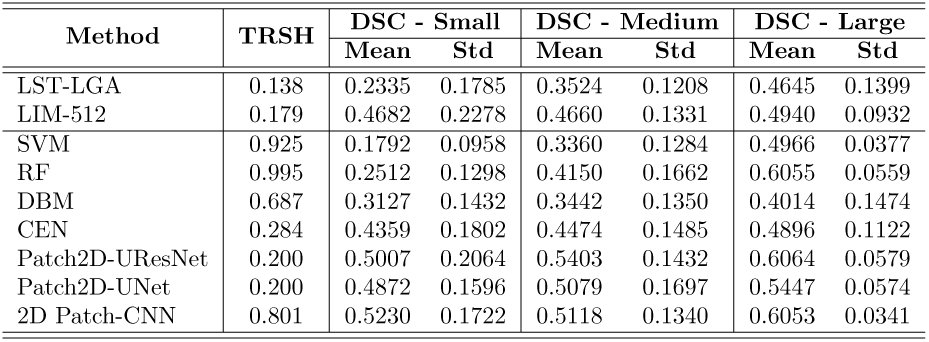
Mean and standard deviation values of dice similarity coefficient (DSC) score’s distribution for all methods tested in this study in respect to WMH burden of each patient (see Table 3). Note that LOTS-IM-512 is listed as LIM-512 in this table.

**Figure 7:**
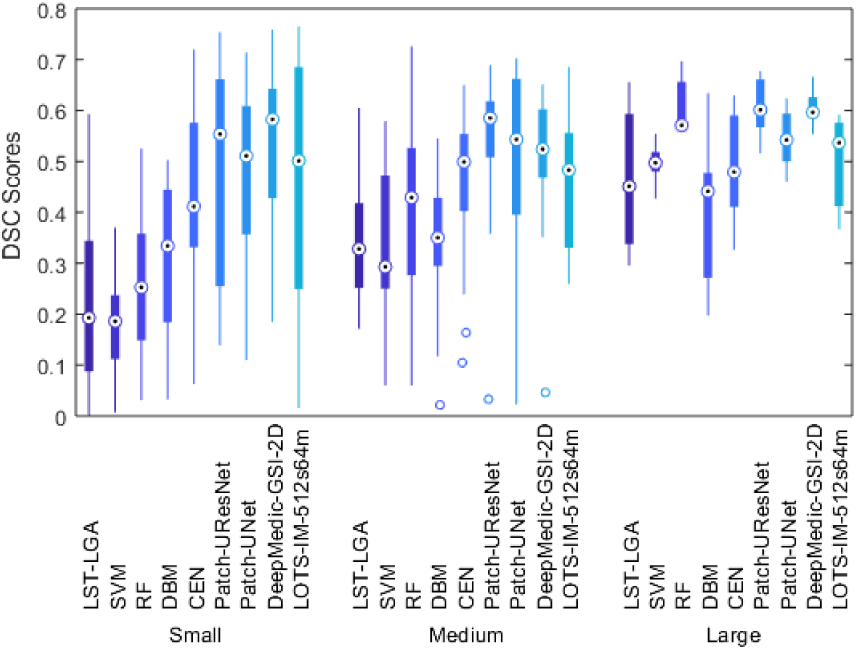
Distributions of dice similarity coefficient (DSC) scores for all methods tested in this study in respect to WMH burden of each patient (see Table 3).

From Figure 7, it can be appreciated that LOTS-IM-512 performed better than LST-LGA in all groups. LOTS-IM-512 also performed better than the conventional supervised machine learning algorithms (i.e. SVM and RF) in “Small” and “Medium” WMH burden groups. Whereas, LOTS-IM-512’s performance was at the level, if not better, than the supervised deep learning algorithms DBM and CEN. However, LOTS-IM-512 still could not beat the *state-of-the-art* supervised deep learning algorithms in any group.

To make this observation clearer, Table 4 lists the mean and standard deviation values that correspond to the box-plot shown in Figure 7. From both Figure 7 and Table 4, it can be observed that the standard deviation of LOTS-IM’s performances in “Small” WMH burden is still relatively high compared to one from the other methods evaluated. However, LOTS-IM’s performance is more stable in “Medium” and “Large” WMH burdens.

#### 7.1.6. Analysis on LOTS-IM’s random sampling

To automatically detect T2-FLAIR’s irregular textures (i.e., WMH) without any expert supervision, LOTS-IM works on the assumption that normal brain tissue is predominant compared with the extent of abnormalities. Due to this assumption, random sampling is used in the computation of LOTS-IM to choose the target patches. Also, note that by using random sampling, the distribution of sampled target patches will follow the underlying distribution of brain tissues in a particular slice. However, it raises an important question on the stability of LOTS-IM’s performance to produce the same level of results for one exact MRI data, especially using different number of target patches. In the first experiment, we randomly chose one MRI data out of the 60 MRI data that we have and ran LOTS-IM 10 times using different number of target patches. Each result was then compared to the ground truth and listed in Table 5. From this experiment, we can see that each setting produced low standard deviation values which indicates that the results are closely distributed around the corresponding mean values. However, there is an indication that higher deviations are produced when using fewer number of target patches.

**Table 5:** Distribution metrics (mean and standard deviation) based on DSC for each LOTS-IM’s settings. Each LOTS-IM setting is tested on a random MRI data 10 times.

| No | Method | TRSH | DSC |  |
| --- | --- | --- | --- | --- |
|  |  |  | mean | std |
| 1 | LOTS-IM-2048 | 0.178 | 0.5681 | 0.0041 |
| 2 | LOTS-IM-1024 | 0.178 | 0.5901 | 0.0018 |
| 3 | LOTS-IM-512 | 0.178 | 0.5922 | 0.0033 |
| 4 | LOTS-IM-256 | 0.178 | 0.5925 | 0.0075 |
| 5 | LOTS-IM-128 | 0.178 | 0.5848 | 0.0092 |
| 6 | LOTS-IM-64 | 0.178 | 0.5852 | 0.0141 |

In the second experiment, we chose three random MRI data from each group of WMH burden (i.e., “Small”, “Medium” and “Large”) based on Table 3, ran LOTS-IM-512 10 times, and compared the results with the ground truth. The results are listed in Table 6. Similar to the first experiment, low standard deviation values were produced for each subject, regardless of the WMH burden.

**Table 6:** Distribution metrics (mean and standard deviation) based on DSC for subject with different WMH burden. Each subject is tested 10 times using LOTS-IM-512.

| WMH Burden | Subject | DSC |  |
| --- | --- | --- | --- |
|  |  | mean | std |
| “Small” | S1 | 0.2481 | 0.0148 |
|  | S2 | 0.1998 | 0.0038 |
|  | S3 | 0.5516 | 0.0067 |
| “Medium” | S4 | 0.6301 | 0.0058 |
|  | S5 | 0.3044 | 0.0013 |
|  | S6 | 0.2907 | 0.0039 |
| “Large” | S7 | 0.5659 | 0.0037 |
|  | S8 | 0.3623 | 0.0045 |
|  | S9 | 0.5671 | 0.0051 |

The two experiments done for this analysis indicate that LOTS-IM produces stable results of WMH segmentation in multiple test instances regardless of WMH burden while employing a simple random sampling scheme. However, of course, more sophisticated sampling method could be applied to make sure patches of normal brain tissue are more likely to be sampled.

#### 7.1.7. Longitudinal test on MCI/AD patients

In this experiment, we evaluated spatial agreement between the produced results in three consecutive years. For each subject, we aligned Year-2 (Y2) and Year-3 (Y3) MRI to the Year-1 (Y1) using niftyReg through TractoR (Clayden et al., 2011), performed subtraction between the aligned WMH labels of baseline/previous year and follow-up year(s) (i.e., Y2-Y1, Y3-Y2, and Y3-Y1), and then labelled each voxel as either “Grow”, “Shrink” or “Stay”. The voxel is labelled “Grow” if it has value above zero after subtraction, labelled “Shrink” if it has value below zero after subtraction, and labelled “Stay” if it has value of zero after subtraction and one before subtraction. This way, we can see whether the method captures the progression of WMH across time.

Figure 8 depicts the results of longitudinal test listed in Table 7 for all methods (i.e., LST-LGA, LOTS-IM-512, Patch2D-UNet, Patch2D-UResNet and DeepMedic-GSI-2D). In this longitudinal test, LOTS-IM-512 is the second-best performer (underlined) on “Grow”, “Shrink” and “Stay” regions segmentation task evaluated using DSC metric after Patch2D-UResNet (written in bold). This, again, confirms that the LOTS-IM shows comparable performance with the *state-of-the-art* supervised deep learning methods (i.e., Patch2D-UNet, Patch2D-UResNet and DeepMedic-GSI-2D).

**Table 7:** Mean and standard deviation values produced in longitudinal test (see Figure 8). LOTS-IM-GPU-512 is listed as LIM-512 in this table. The best values are written in **bold** while the second-best values are <u>underlined</u>. In this longitudinal test, LIM-512 outperformed LST-LGA while competed with the supervised deep learning methods.

| Method | Dice Similarity Coefficient (DSC) |  |  |  |  |  |
| --- | --- | --- | --- | --- | --- | --- |
|  | Grow |  | Stay |  | Shrink |  |
|  | Mean | Std | Mean | Std | Mean | Std |
| LST-LGA | 0.1301 | 0.0350 | 0.2343 | 0.0199 | 0.2706 | 0.0058 |
| LIM-512 | <u>0.2260</u> | 0.0084 | <u>0.4585</u> | 0.0104 | <u>0.3715</u> | 0.0018 |
| Patch2D-UNet | 0.2242 | 0.0125 | 0.4207 | 0.0125 | 0.3675 | 0.0242 |
| Patch2D-UResNet | **0.2523** | 0.0199 | **0.4664** | 0.0211 | **0.3912** | 0.0044 |
| 2D Patch2DCNN | 0.1440 | 0.0228 | 0.4066 | 0.0298 | 0.3660 | 0.0129 |

**Figure 8:**
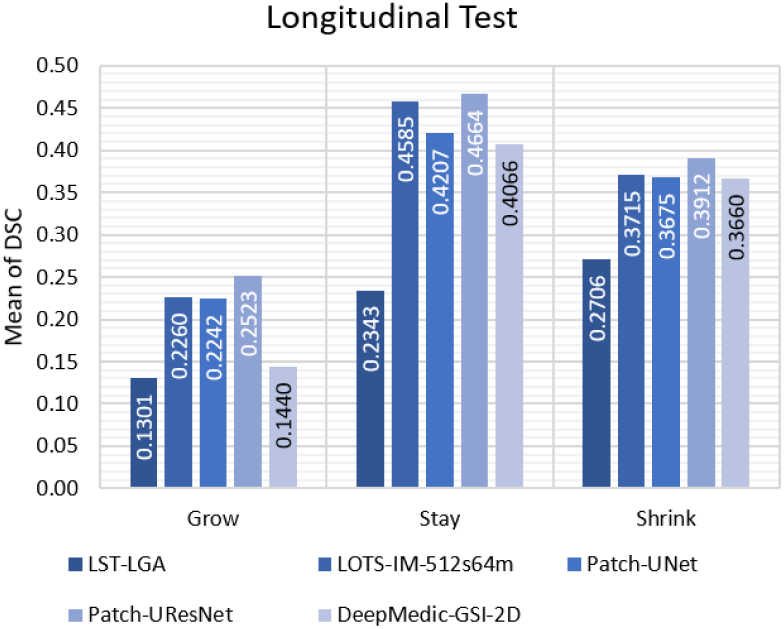
Quality of spatial agreement (mean of DSC) of the produced results in longitudinal test. Longitudinal test is done to see the performance of tested methods in longitudinal data set of MRI (see Table 7 for full report).

#### 7.1.8. Correlation with visual scores

In this experiment, we want to see how close IAM’s results correlate with visual rating scores of WMH, specifically Fazekas (Fazekas et al., 1987) and Longstreth’s visual scores (Longstreth et al., 1996).

We calculated Spearman’s correlation coefficient 1) between the total Fazekas score (i.e., the sum of periventricular and deep white matter hyper-intensities) and manual/automatic WMH volumes and 2) between Longstreth’s total score and manual/automatic WMH volumes. The results are listed in Table 8.

**Table 8:** Non-parametric correlation using Spearman’s correlation coefficient between WMH volume and Fazekas and Longstreth visual ratings.

| Visual Rating<br>Method | Fazekas (Total) |  | Longstreth |  |
| --- | --- | --- | --- | --- |
|  | Spearman’s Corr. |  | Spearman’s Corr. |  |
| | $\rho$ | P | $\rho$ | P |
| Manual label | 0.7562 | $1.04 \times 10^{-12}$ | 0.7752 | $1.45 \times 10^{-12}$ |
| LST-LGA | 0.5718 | $3.38 \times 10^{-12}$ | 0.4813 | $1.50 \times 10^{-4}$ |
| LIM-2048 | 0.4727 | $2.05 \times 10^{-4}$ | 0.4579 | $3.42 \times 10^{-4}$ |
| LIM-1024 | 0.4892 | $1.13 \times 10^{-4}$ | 0.4849 | $1.32 \times 10^{-4}$ |
| LIM-512 | 0.5010 | $7.19 \times 10^{-5}$ | 0.5065 | $5.82 \times 10^{-4}$ |
| LIM-256 | 0.5009 | $7.22 \times 10^{-5}$ | 0.5085 | $5.37 \times 10^{-4}$ |
| LIM-128 | 0.4505 | $4.38 \times 10^{-4}$ | 0.4946 | $9.22 \times 10^{-4}$ |
| LIM-64 | 0.4393 | $6.30 \times 10^{-4}$ | 0.4858 | $1.28 \times 10^{-4}$ |
| SVM | 0.4062 | $1.70 \times 10^{-2}$ | 0.3602 | $5.90 \times 10^{-3}$ |
| RF | 0.2447 | $6.66 \times 10^{-2}$ | 0.2128 | $1.12 \times 10^{-1}$ |
| DBM | 0.2436 | $6.79 \times 10^{-2}$ | 0.1659 | $2.17 \times 10^{-1}$ |
| CEN | 0.2359 | $7.74 \times 10^{-2}$ | 0.3618 | $5.70 \times 10^{-3}$ |
| Patch2D-UResNet | 0.3602 | $5.90 \times 10^{-3}$ | 0.5171 | $3.80 \times 10^{-5}$ |
| Patch2D-UNet | 0.4618 | $2.99 \times 10^{-4}$ | 0.5140 | $4.33 \times 10^{-5}$ |
| DeepMedic-GSI-2D | 0.7054 | $9.01 \times 10^{-10}$ | 0.8664 | $3.19 \times 10^{-18}$ |

Table 8 shows that, although not much better, all LOTS-IM methods are highly correlated with visual rating clinical scores. It is also worth to mention that LST-LGA produced WMH segmentation results which are highly correlated with visual ratings but produced the lowest DSC metric of all (see Table 1). On the other hand, LOTS-IM produced high values of DSC metric and high correlation with visual scores at the same time. Visual inspection of the LOTS-IM results revealed systematic false positive detection in the cerebellum, aqueduct, Sylvian fissure and some cortical regions. These errors are consistent with those reported by other WMH segmentation methods (Hernández et al., 2010).

### 7.2. Results on MS Patients

In this section, LOTS-IM’s performance for MS lesion segmentation and analysis of progression are presented and discussed.

#### 7.2.1 LOTS-IM for MS lesion segmentation

In this experiment, LOTS-IM-512, LOTS-IM-256, LOTS-IM-128 and LOTS-IM-64 were chosen to represent LOTS-IM as they are comparable with LST-LGA in terms of speed while having no significantly different performance to LOTS-IM-1024 and LOTS-IM-2048. Furthermore, unlike in previous experiments, only the Dice similarity coefficient (DSC) metric and the two-sided Wilcoxon signed rank test were used to evaluate LOTS-IM’s performance for MS lesion segmentation.

Table 9 shows the performance of LOTS-IM and LST-LGA for MS lesion segmentation in the dataset 2. The table provides both results for the whole data set (i.e. 30 MRI scans from the UMCL) and in the three groups of MS lesions burden (i.e., “Small”, “Medium” and “Large” based on Table 10).

**Table 9:** Experiment results produced by LST-LGA and LOTS-IM on MS patients from the data set 2. The results are based on the Dice similarity coefficient (DSC) where the best value is <u>underlined</u>. Two-sided Wilcoxon signed rank test is performed between LST-LGA and different settings of LOTS-IM to determine whether the performance of these two methods significantly differ from one another or not **Abbreviations**: “TRSH” for optimum threshold while “h” and “p” for Wilcoxon’s output values.

| Method | TRSH | DSC - All Scans |  |  | Speed (min) | DSC - Small |  |  | DSC - Medium |  |  | DSC - Large |  |  |
| --- | --- | --- | --- | --- | --- | --- | --- | --- | --- | --- | --- | --- | --- | --- |
|  |  | mean (std) | h | p |  | mean | h | p | mean | h | p | mean | h | p |
| LST-LGA | 0.564 | <u>0.5145</u> (0.241) | - | - | 45.12 | <u>0.1651</u> | - | - | 0.5131 | - | - | <u>0.6793</u> | - | - |
| LOTS-IM-512 | 0.128 | 0.4941 (0.248) | 0 | 0.4174 | 30.44 | 0.1386 | 0 | 0.6406 | 0.5400 | 0 | 1.0000 | 0.6506 | 0 | 0.5540 |
| LOTS-IM-256 | 0.138 | 0.4935 (0.248) | 0 | 0.3812 | 14.29 | 0.1386 | 0 | 0.3828 | <u>0.5437</u> | 0 | 1.0000 | 0.6486 | 0 | 0.5540 |
| LOTS-IM-128 | 0.148 | 0.4908 (0.251) | 0 | 0.3147 | 9.54 | 0.1317 | 0 | 0.4609 | 0.5266 | 0 | 0.8750 | 0.6514 | 0 | 0.5540 |
| LOTS-IM-64 | 0.168 | 0.4842 (0.254) | 0 | 0.1631 | 9.35 | 0.1211 | 0 | 0.1484 | 0.5165 | 0 | 0.8750 | 0.6476 | 0 | 0.5528 |

**Table 10:**
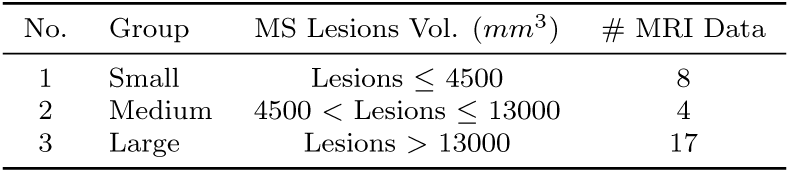
Three groups of MS patients based on MS lesions load.

As Table 9 shows, LST-LGA performed better than LOTS-IM methods in terms of DSC metric. However, the differences between both methods’ performance were not significant as per the two-sided Wilcoxon signed rank test with 5% significance level. Similar results are observed in MS lesion segmentation per MS lesion burden where the differences between these two methods’ performances are small in term of DSC metric and not significant in all groups. Thus, we can say that LOTS-IM performs similar to LST-LGA for MS lesion segmentation in this data set. However, LOTS-IM is more flexible than LST-LGA in term of speed as LOTS-IM can be set to run faster than LST-LGA by using less number of target patches while having small to none degradation in performance.

Furthermore, it is also worth to mention that this experiment shows that LOTS-IM is more robust than LST-LGA when applied to different data sets. Note that ADNI data set has smaller resolution than MS data set and the T2-FLAIR hyperintensities are mainly thought to be from different aetiologies (i.e., WMH of presumable vascular origin vs MS lesions). While LOTS-IM performed stable without big differences on both ADNI and MS data sets, LST-LGA’s performance dropped significantly in the ADNI data set.

#### 7.2.2. Applicability to assess MS lesion progression

The evolution of interval or enlarging lesions on T2-dependent imaging are key criterion for assessing MS disease activity, which informs clinical decision-making and as a surrogate endpoint for clinical trials of therapeutic agents. We coregistered the raw (i.e. not post-processed) LOTS-IM output obtained from the baseline and follow-up FLAIR images from the data set 3, to a mid-space and subtracted both maps. Then, we performed 3d connected component analysis to the “positive” and “negative” regions of the subtracted maps, being these regions comprised by the voxels with modular values higher than 0.18. We followed the same thresholding criterion as the one followed to extract the WMH to neglect subtle differences due to misregistrations, cortical effects, or differences in image contrast not related to the disease. We counted the “positive” spatial clusters (i.e. connected components) with IM values (i.e. in at least one voxel) higher than 0.7 and labelled those spatial clusters as “New or enlarged lesions”. We summed the areas (in number of voxels) covered by all the “positive” and “negative” spatial clusters weighted by their mean IM value, separately and subtracted them to determine the overall change and rated it in none-low (less than 25% of the “positive” areas), low-moderate (between 25 and 50% of the “positive” areas) and moderate-high (more than 50% of the “positive” areas. Two expert raters, independently and blind to any quantitative analysis, visually assessed the baseline and follow-up images and identified the number of new and/or enlarged MS lesions, and rated the disease progression. Discrepancies among raters were discussed and a final result was agreed. Table 11 shows the results from both assessments.

**Table 11:** Visual expert vs. LOTS-IM longitudinal MS lesion assessments in 10 treatment-free MS patients (data set 3)

| Patient | Visual expert assessment |  | LOTS-IM output |  |
| --- | --- | --- | --- | --- |
|  | New or enlarged lesions | Dis. progres. | New or enlarged lesions | Dis. progres. |
| 1 | 0 | None-Low | 0 | None-Low |
| 2 | 1 | None-Low | 3 | Moderate-High |
| 3 | 0 | None-Low | 2 | None-Low |
| 4 | 1 | Moderate-High | 5 | Low-Moderate |
| 5 | 0 | None-Low | 1 | None-Low |
| 6 | 2 | Moderate-High | 3 | Moderate-High |
| 7 | 5 | Moderate-High | 6 | Low-Moderate |
| 8 | 3 | Moderate-High | 1 | None-Low |
| 9 | 2 | Moderate-High | 5 | Moderate-High |
| 10 | 6 | Moderate-High | 2 | Moderate-High |

The agreement between LOTS-IM and the expert assessment in rating the disease progression was 80%. The criterion followed to count the new or enlarged lesions from the LOTS-IM needs to be revised though, as the automatic count does not reflect actual disease change in some cases. MS lesions which are active at one imaging time point and subsequently quiescent frequently decrease in size; summed volume measures may therefore not identify active disease when some lesions are enlarging and others are shrinking. The automatic processing of LOTS-IM output as described above, identified changes produced by CSF flow artifacts in the choroid plexus, temporal poles and junction between the septum and the callosal genu or splenium (Figure 9). These, although genuine signal changes, are not related to the disease. Instead, these are particular confounders in MS where genuine lesions frequently abut ventricular margins. In addition, the centre of T1-weighted “black holes”, not hyperintense in baseline FLAIR images but hyperintense in the follow-up FLAIR, which maybe due to variation in efficacy of fluid signal suppression, was also counted as new lesion (Figure 9). Although initial results are promising, post-processing to correct these “false” positives and negatives will be necessary for applying LOTS-IM to clinical research in MS.

**Figure 9:**
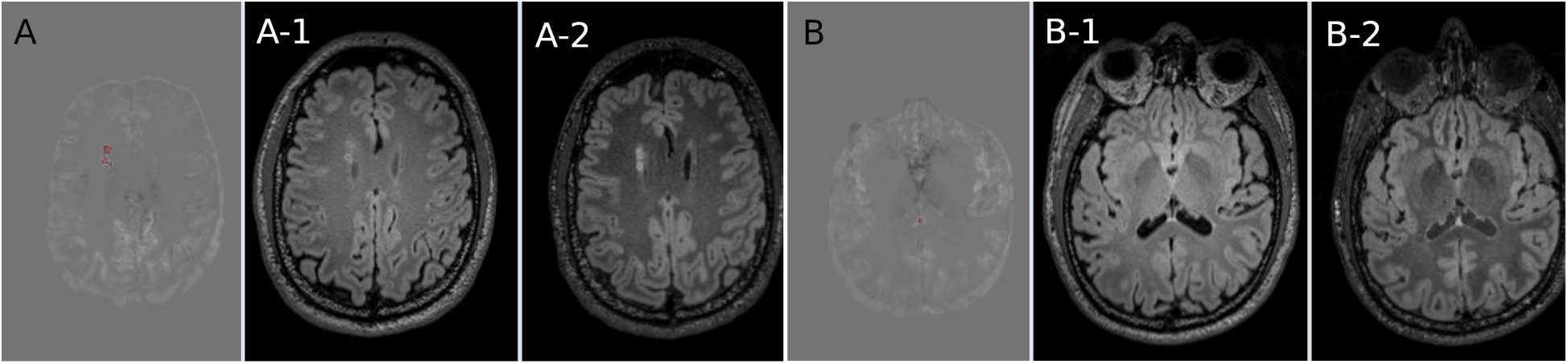
Two examples (i.e. A and B) of WMH change captured by LOTS-IM that do not represent actual disease progression from the neuroradiological perspective. A-1 and B-1 are the baseline FLAIR axial slices and A-2 and B-2 are the corresponding follow-up FLAIR slices. The “new” lesions captured by LOTS-IM are represented in red on the “change” LOTS-IM. In A, the two “new” lesions are void regions on the baseline FLAIR, in the centre of two neighbouring hyperintensities, which are hyperintense at follow-up. In B, the “new” lesion is in reality a CSF flow artefact in the intersection between the septum and the splenium of the corpus callosum.

## 8. Conclusion and Future Work

In this study, we have presented the use of LOTS-IM for WMH segmentation, MS lesion segmentation and analysis of MS lesion progression. We have also shown that the optimisation of the irregularity map method presented (LOTS-IM) accelerates processing time by large margin without excessive quality degradation compared with the previous iterations (IAM and OTS-IAM). LOTS-IM speeds up the overall computation time, attributable not only to implementation on GPU, but also to the use of a limited number of target patch samples. In addition, we have evaluated LOTS-IM in different scenarios, which was not done in previous studies.

Unlike other WMH segmentation methods, LOTS-IM successfully identifies and represents both non-WMH regions and WMH regions using irregularity map, including the “penumbra” of WMH. Despite not being a WMH segmentation method *per se*, LOTS-IM can be applied for this purpose by thresholding the value of the irregularity map. Being unsupervised confers an additional value to this fully automatic method as it does not depend on expert-labelled data, and therefore is independent from any subjectivity and inconsistency from human experts, which typically influence supervised machine learning algorithms. Our results show that LOTS-IM outperforms LST-LGA, the current *state-of-the-art* unsupervised method for WMH segmentation, conventional supervised machine learning algorithms (SVM and RF), and some supervised deep learning algorithms (DBM and CEN). Our results also show that LOTS-IM has comparable performance with the *state-of-the-art* supervised deep learning algorithms DeepMedic, UResNet, and UNet.

On MS lesion segmentation, LOTS-IM’s performance is similar to the LST-LGA. While LOTS-IM could not outperform LST-LGA on MS lesion segmentation, LOTS-IM is more flexible than LST-LGA as its computation speed can be accelerated by using less target patches. LOTS-IM is also more robust than LST-LGA as it performed stable (i.e., without any big difference) in both ADNI and MS patient data sets. This contrasts to LST-LGA where its performance dropped significantly on ADNI data set when dealing with WMH segmentation. Furthermore, LOTS-IM also performed well in assessing MS lesion progression where the agreement between LOTS-IM and experts’ visual assessment was 80%.

One limitation of LOTS-IM is the influence that the quality of brain masks (i.e., CSF and NAWM) has in its performance. We have shown that random sampling has a small impact to the final result on WMH segmentation, but more sophisticated sampling could be used as well. Some improvements also could be done by adding or using different sets of brain tissues masks other than CSF and NAWM, such as cortical and cerebrum brain masks.

We believe that the irregularity map could provide unsupervised information for pre-training supervised deep learning, such as UResNet and UNet. In (Rachmadi et al., 2018a), UNet successfully learned the irregularity map produced by LOTS-IM. Progression/regression of brain abnormalities also can be achieved with LOTS-IM(Rachmadi et al., 2018a). Due to its principle, it could be applicable to segment brain lesions in CT scans or different brain pathologies, but further evaluation would be necessary. Further works could also explore its implementation on a multispectral approach that combines different MRI sequences. The implementation of LOTS-IM on GPU is publicly available on the following GitHub page^8^

## Acknowledgements

The first author would like to thank Indonesia Endowment Fund for Education (LPDP) of Ministry of Finance, Republic of Indonesia, for funding his study at School of Informatics, the University of Edinburgh. Funds from Row Fogo Charitable Trust (Grant No. BRO-D.FID3668413)(MCVH) are also gratefully acknowledged.

Data collection and sharing for this project was partially funded by the Alzheimer’s Disease Neuroimaging Initiative (ADNI) (National Institutes of Health Grant U01 AG024904) and DOD ADNI (Department of Defense award number W81XWH-12-2-0012). ADNI is funded by the National Institute on Aging, the National Institute of Biomedical Imaging and Bioengineering, and through generous contributions from the following: AbbVie, Alzheimers Association; Alzheimers Drug Discovery Foundation; Araclon Biotech; BioClinica, Inc.; Biogen; Bristol-Myers Squibb Company; CereSpir, Inc.; Cogstate; Eisai Inc.; Elan Pharmaceuticals, Inc.; Eli Lilly and Company; EuroImmun; F. Hoffmann-La Roche Ltd and its affiliated company Genentech, Inc.; Fujirebio; GE Healthcare; IXICO Ltd.; Janssen Alzheimer Immunotherapy Research and Development, LLC.; Johnson and Johnson Pharmaceutical Research and Development LLC.; Lumosity; Lundbeck; Merck and Co., Inc.; Meso Scale Diagnostics, LLC.; NeuroRx Research; Neurotrack Technologies; Novartis Pharmaceuticals Corporation; Pfizer Inc.; Piramal Imaging; Servier; Takeda Pharmaceutical Company; and Transition Therapeutics. The Canadian Institutes of Health Research is providing funds to support ADNI clinical sites in Canada. Private sector contributions are facilitated by the Foundation for the National Institutes of Health (www.fnih.org). The grantee organization is the Northern California Institute for Research and Education, and the study is coordinated by the Alzheimers Therapeutic Research Institute at the University of Southern California. ADNI data are disseminated by the Laboratory for Neuro Imaging at the University of Southern California.

FutureMS is supported by an exemplar grant from the Stratified Medicine Scotland Innovation Centre and funding from Biogen, Inc (Cambridge, Massachusetts, https://www.biogen.com/). RM (and partially MCVH) salaries are supported by the CSO-PME grant to the Stratified Medicine Scotland Innovation Centre.

## Footnotes

1 https://sourceforge.net/projects/bric1936/files/MATLAB_R2015a_to_R2017b/BRIClib/

2 https://surfer.nmr.mgh.harvard.edu/

3 http://adni.loni.usc.edu/

4 http://hdl.handle.net/10283/2214

5 http://adni.loni.usc.edu/wp-content/uploads/how_to_apply/ADNI_Acknowledgement_List.pdf

6 http://lit.fe.uni-lj.si/tools

7 https://future-ms.org/

8 https://github.com/febrianrachmadi/lots-iam-gpu.

